# Protein as evolvable functionally-constrained amorphous matter

**DOI:** 10.1101/2022.07.17.500335

**Authors:** Madhusmita Tripathy, Anand Srivastava, Srikanth Sastry, Madan Rao

## Abstract

We explore current ideas around the representation of a protein as an amorphous material, in turn represented by an abstract graph 𝒢 with edges weighted by elastic stiffnesses. By embedding this graph in physical space, we can map every graph to a spectrum of conformational fluctuations and responses (as a result of say, ligand-binding). This sets up a “genotype-phenotype” map, which we use to evolve the amorphous material to select for fitness. Using this, we study the emergence of allosteric interaction, hinge joint, crack formation and a slide bolt in functional proteins such as Adenylate kinase, HSP90, Calmodulin and GPCR proteins. We find that these emergent features are associated with specific geometries and mode spectra of floppy or liquid-like regions. Our analysis provides insight into understanding the architectural demands on a protein that enable a prescribed function and its stability to mutations.

## INTRODUCTION

Only a small fraction of the allowable protein ‘universe’ constitutes real biological proteins [1, 2]. For example, of the 20^300^ number of possible sequences of a polypeptide chain with ∼ 300 residues that can potentially be generated from the naturally available 20 amino acids, living systems such as *Saccharomyces cerevisiae* exhibit only ∼ 10^4^ of these [2–4]. This *dimensional reduction* comes about because, out of the numerous possible proteins, only a small subset are functionally relevant, robust and explored by evolution. We have, for many years, been interested in understanding the architectural demands on a protein that enable a specific function, and its stability to mutations, fluctuations and cycles of performance. Some aspects of this program are not new and recently, rather elegant theoretical formalisms have emerged [5–9]. Here we offer our perspective on this problem.

We focus on proteins that undergo significant conformational changes between their native and functional states. We first consider ‘allosteric proteins’, where the intriguing mechanism of ‘action-at-a-distance’ drives function. Motivated by the tantalising similarities between functional proteins and amorphous materials, in terms of molecular packing [10], free energy landscape [11] and relaxation mechanisms [12], we explore if allosteric regulation proceeds via emergence of ‘allosteric chains’, reminiscent of ‘force chains’ in granular media [13]. There are two proposed mechanisms of allostery – the *induced-fit* mechanism, where the conformational switch depends on a ligand induced change in protein conformation that leads to specificity of enzyme action [14], and the *conformational selection* mechanism, where the enzyme explores a multiplicity of conformation states, independent of ligand structure and occupancy, which are then differentially stabilized by the ligand [15, 16]. Since allosteric propagation and binding scenarios in proteins span a repertoire of selection and adjustment processes, it is likely that both these mechanisms could be operative in the same protein in physiological settings [17–20]. Here we focus on induced fit proteins, such as Adenylate and Guanylate kinase [21–25], HSP90 [26], Calmodulin [27–29] and GPCR proteins [30–32], and ask what are the necessary physical (architectural) features that the protein must have in order to perform a specific function with high fidelity.

To do this, we need a coarse-grained representation of a protein that is appropriate for this task. A protein represented as a heteropolymer [33] is indeed a convenient starting point if the question pertains to the dynamics of folding into a native state, or to the dynamics of assembly driven by multivalent interactions of intrinsically disordered proteins [34]. However, a coarse-grained description of changes in protein conformation in the native state, either as a result of spontaneous fluctuations or induced by ligand binding, or during the process of chemical reaction, requires a different starting point. We need a representation that enables a classification of the low energy excitations and modes of deformation about the native state of a protein [24]. This would involve accounting for inter-monomer (or inter-sector) [35, 36] interactions of varying strengths, both along the heteropolymer backbone and across it, giving it a three-dimensional character. This suggests that the appropriate coarse-grained description for deformations of a functional protein is to treat the protein as a 3-dimensional amorphous solid with heterogeneous interactions that have been designed to facilitate a prescribed function with high fidelity. The strategy that we will use to *design* the heterogeneous interactions is akin in spirit to a *gain of function* approach [37, 38]. The ability to render a specific function with high fidelity puts constraints on the free energy landscape explored by the amorphous solid.

## I. REPRESENTATION OF A PROTEIN AS AN AMORPHOUS MATERIAL

Here we make precise the representation of a protein as an amorphous solid. For simplicity, we will consider proteins that have a large molecular weight and are globular, with a well defined “bulk” and “surface”. A globular protein is a linear heteropolymer with side groups, which in its native conformation is folded up in a ball. This enables each monomer to interact with the rest of the monomers across 3d space, via interactions of varying bond strengths. It is in this setting that we define the genotype-phenotype space and the representation as an amorphous solid.

### A. “Genotype” Space

Let the set of *amino acids* (monomer types) be {*A*_*i*_} : *i* = 1, … *K*, each characterised by a hydrodynamic radius {*a*_*i*_} and the set of *bond* types be {*B*_*α*_} : *α* = 1, … *M*, with *M* ≪ *K*, each characterised by a bond-strength {*b*_*α*_} (Fig. 1(i)). A realisation of a “protein” is a weighted graph 𝒢 = {𝒱, ℰ}, with the vertices 𝒱 taken from {*A*_*i*_} and edges ℰ taken from {*B*_*α*_}. Note that a given vertex can have any number of edges emanating from it; it is greater than 1 (if surface vertex) or 2 (if bulk vertex) and less than a maximum *E*_*max*_.

**FIG. 1.**
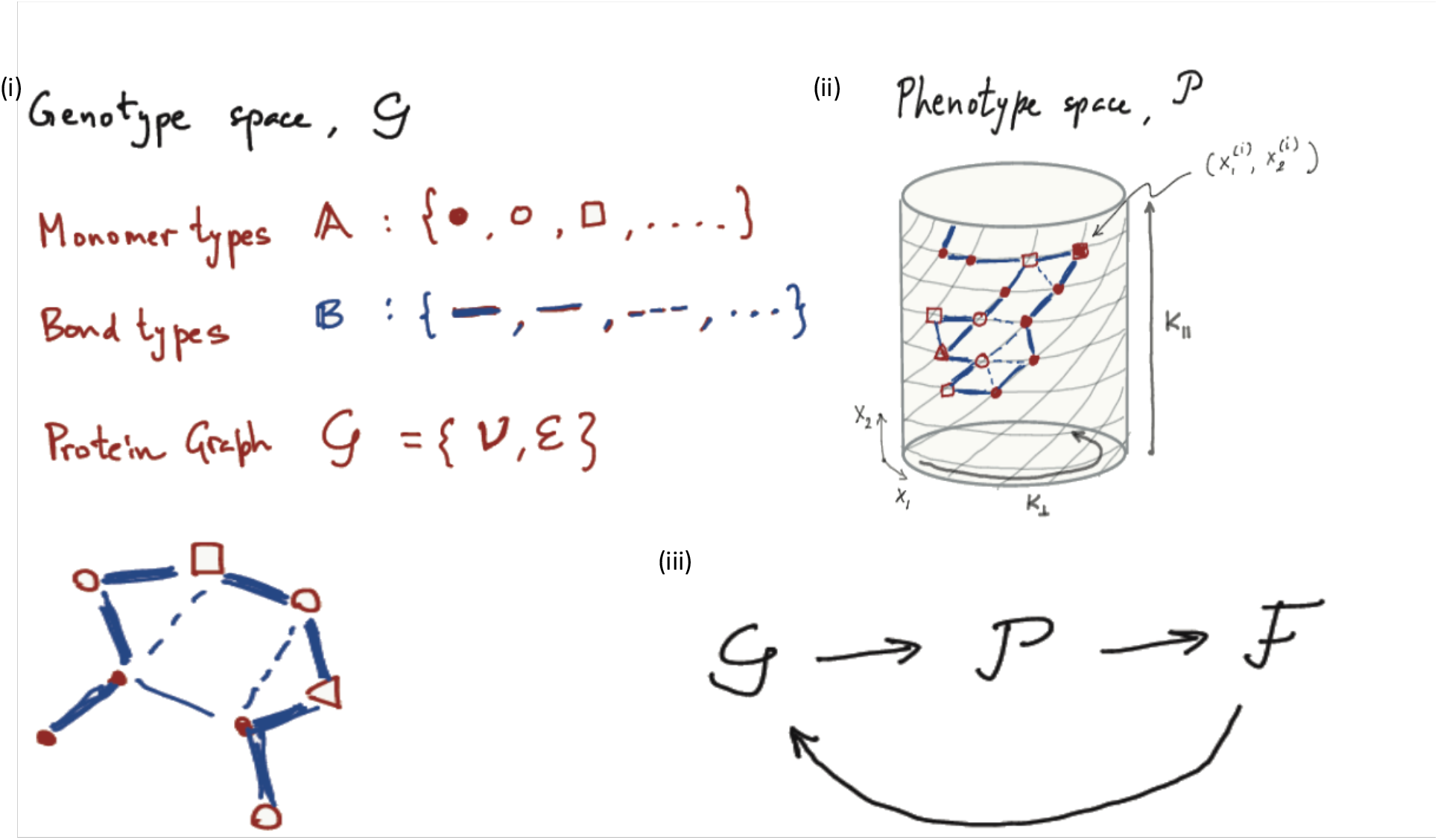
(i) *Genotype* space 𝒢 constructed from the set of monomers and bonds with different stiffnesses to generate a weighted protein graph 𝒢 = {𝒱, ℰ} representing the abstract protein network. (ii) *Phenotype* space 𝒫 obtained by embedding the graph 𝒢 in physical space. This embedding assigns coordinates to the vertices. We draw it on a cylinder to depict that we impose periodic boundary conditions in the *x*_1_-direction and free boundaries in the *x*_2_-direction. This choice of boundary conditions is dictated by the nature of the fitness function ℱ. (iii) Every network in 𝒢, gets embedded in 𝒫 from which we compute the fitness function ℱ. We then change the network in 𝒢 and iterate till we reach the optimum fitness.

Together these constitute the *genotype* space 𝒢.

### B. “Phenotype” Space – Embedding in physical space

As shown in Fig. 1(ii), we embed this graph 𝒢 in *physical space*, that is to say, the *N* vertices are embedded in Euclidean space of *d*-dimensions ℝ _*d*_ (with coordinates 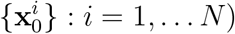).

With this embedding, each vertex is subjected to forces arising from steric repulsion upon contact and short-range harmonic extensional springs from the connecting bonds. In addition, one could include contributions to the force, such as bending and torsion. This sets the stage for viewing the protein as an amorphous solid with heterogeneous spring constants.

Because the protein is a polymer with a defined backbone characterised by stronger peptide bonds, the energy scales associated with the extensional springs in the above representation will show a clear separation in bond strengths. We will refer to the peptide bonds of the backbone as *strong* bonds and the interactions such as electrostatic, hydrophobic, hydrogen bonding, disulfide and salt bridges, van der Waals, collectively as *weak* bonds. Neighbouring monomers that do not interact will be connected by a *non-bonding* edge. Given that the protein is a linear polymer, every bulk vertex will have two strong bonds emanating from it.

Together these define the *phenotype* space 𝒫.

### C. Fidelity of function as Fitness

Having established the Genotype-Phenotype map 𝒢 → 𝒫, we would like to drive changes in the genotype space to arrive at a desired phenotype. We do this by defining a *fitness function*.

Since we will be concerned with native proteins that undergo specific conformational change in response to a local external stimulus, such as ligand binding, the fitness function must describe the fidelity and specificity of the conformational change. Thus, in general, we define fitness as a scalar function of the displacements of the vertices of the physical graph, i.e. ℱ : 𝒫 → ℝ. This function has as *input* ℐ, the prescribed displacement vectors of a subset of vertices *i* ∈ ℐ ⊂ 𝒫, and as *output* 𝒪, a scalar function of the displacement vectors of a different subset of vertices *j* ∈ 𝒪 ⊂ 𝒫. The goal is to sample the genotype space 𝒢 and optimise the fitness function ℱ over the space of phenotypes 𝒫. In Sect. II we will consider several examples of this fitness function ℱ.

### D. Optimisation algorithm

While our proposed optimisation algorithm should hold in any dimension, we will for convenience, describe the procedure in two spatial dimensions. We start with a phenotype graph 𝒫 with vertices on a triangular lattice of dimension *K*_∥_ × *K*_⊥_ (Fig. 1(ii)), and edges connecting nearest neighbour vertices, with periodic boundary conditions. Let the initial coordinates of the vertices be 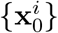.

For the problem at hand, we can, without loss of generality, take all the monomers to be the same and assign all the genotypic diversity to the bonds. Thus, we randomly assign the weight of an edge to be {*b*_*α*_} : *α* = 1, … *M* with probability *p*_*α*_, where *b*_1_ ≡ 0 corresponds to the non-bonded edges. A useful parameter in the model is the number fraction of bonded edges *ϕ*_0_. This assignment should be subjected to constraints, such as ensuring a polymer backbone, i.e. that there exists one and only one path in 𝒫 comprising strong covalent bonds alone that spans all vertices, but for now we will ignore this constraint.

Given a realisation of bond-strength *b*’s on the phenotype graph 𝒫, we can compute real space displacements **u**^*i*^ of every vertex, by minimising the total elastic energy, 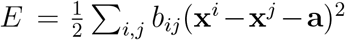 with respect to **u**^*i*^, where 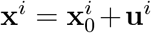 and 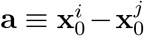. If our physical embedding was associated with a bath of temperature *T*, we could in principle even compute the displacement fluctuations at every vertex. These measurable physical quantities will depend on the spring constants that reside in the bonds and in the hydrodynamic radii that reside in the vertex.

Now for every realisation of bond-strengths {*b*_*ij*_} on the phenotype graph 𝒫, we can compute the fitness function ℱ for the prescribed input. We then change the realisation of {*b*_*ij*_}, and repeat the calculation. By sampling over all the realisations of *b*, we arrive at one that optimises ℱ for the same fixed input. In practice this is hard because the dimensionality of the search space goes as *M* ^*N*^, a very large number. We will therefore restrict the bond strengths to {*b* = 0, 1}, in units of a typical energy scale, and sample the genotype space 𝒢 using a Metropolis Monte Carlo sampling scheme.

We implement the algorithm as follows:

We first prepare the system by distributing the bond strengths {0, 1} randomly, such that with probability *p* the bond strength is 1; this specifies the number fraction of bonded edges *ϕ*_0_. We construct a well-defined, physically motivated, fitness function ℱ (with nice convergence properties). And choose a large *N*, a large enough *p* to ensure percolation and boundary conditions that are either open or periodic. Then,

1. We provide fixed displacement vectors for the input vertices. In response to this localized strain, all bonds with nonzero stiffness will elastically deform. We then compute the displacements {**u**^*i*^} of all the vertices, that minimise the total elastic energy,

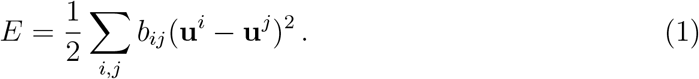
2. Using this energy minimized displacement vectors of the output vertices, we compute the fitness function ℱ. This will be large in general.
3. We now make moves in genotype space 𝒢 (mutations), which corresponds to moving in bond space {*B*_*i*_}. For simplicity, we restrict the space of moves to those that interchange the 0’s and 1’s (bond exchange moves). This fixes the number fraction of bonded edges at its initial value *ϕ*_0_. This is not necessary, one could easily study moves which sample number fractions spread about *ϕ*_0_ (as an aside, altering the value of *ϕ*_0_ can lead us to study issues surrounding isostaticity or overconstrained configurations).
4. We then repeat the calculation and determine the new fitness. We follow this procedure till the fitness ℱ is minimized.

In order to efficiently sample 𝒢 to minimise ℱ, especially when *N* is large, one might choose a simulated annealing scheme, with a fictitious temperature *T*_*f*_. For any nonzero *T*_*f*_, there will be a distribution of optimal configurations, the true optimal network will be obtained by slowly taking *T*_*f*_ → 0.

In practice, we have implemented the above algorithm on a triangular lattice with the number of vertices *N* = 156 arranged in a 12 × 13 grid. We have used a slightly distorted lattice to avoid straight lines of vertices and that result in the appearance of floppy modes [5]. The number of strength-1 bonds, *N*_*S*_ = 360, which we fix throughout the simulation. This in turn fixes the average coordination number, *z* = 2*N*_*S*_*/N* = 5. In addition, the vertices are also connected to their next neighbors via weak springs with stiffness 10^−4^. Periodic boundary condition is imposed based on the specific case being modeled, as specified in Sect. II.

The binding of the ligand is modeled by imposing a displacement field, {**u**^ℐ^}, at the input vertices *i* ∈ ℐ (we take it to be 4 adjacent vertices located at the center of the lower boundary of the grid). Such an imposed displacement results in a deformation of the entire network, leading to a displacement, 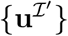, at every other vertex of the network. We numerically evaluate 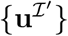 by solving the corresponding global stiffness matrix.

All vertices obey local force balance. Thus for the vertices *i* ∈ ℐ, the external forces required to impose the displacements should balance the internal elastic forces, while for the vertices *j* ∈ ℐ_′_ (the complement of ℐ), the internal elastic forces should add up to zero. In block matrix form,

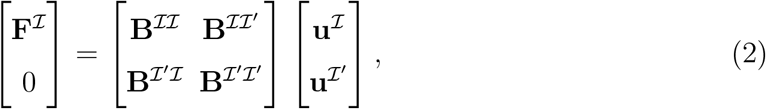

where **B** is the block stiffness matrix. The unknown displacements can be obtained by simple matrix inversion.

Every time we move through the genotype space, we change the topology of the network, and construct a new **B**, which is then used to calculate the unknown displacements 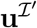. Under this evolution, we search for networks that generate a response, which matches a target displacement, 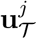, at sites *j* ∈ 𝒪 located far from the input stimulus ℐ. The fitness of the network is evaluated in terms of the deviation of the displacement field at the output sites from its target value,

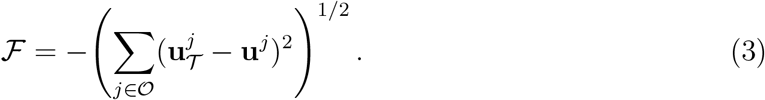

To evolve towards the optimum in this non-convex optimization problem, we perform a Monte Carlo simulation using Metropolis sampling at fictitious temperature *T*_*f*_ = 0.01. The simulation is performed for 5 × 10^5^ steps, where the fitness value usually converges within 100 Monte Carlo steps. We show a movie of the evolution of the network towards optimality in *Network Evolution*.

In the next section, we employ this algorithm to study four different functional proteins. We then characterize the optimised network in terms of the spatial profiles of the mean coordination number and displacement.

## II. EMERGENCE OF FUNCTIONAL PROTEINS

Amongst the quantities we measure are the distributions means and fluctuations of scalars such as (i) averaged local coordination number (number of bonds per site with weight 1), (ii) mean square displacement (SD) at every vertex 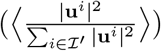. This allows us to classify the variety of protein types according to the relative fraction of liquid to solid regions and geometry of these liquid regions. Using the above genotype-phenotype map, we study the emergence of allosteric interaction, hinge joint, crack formation and a slide bolt in functional proteins, such as Adenylate kinase, HSP90, Calmodulin and so on.

### A. Allosteric proteins with slide bolt behavior

In this case, the active site consists of 4 consecutive vertices on the top boundary. Such a representation models the case of globular allosteric proteins, where the active and allosteric sites are located at specific distant sites, each comprising a small part of the protein surface. In the abstract network, the stimulus site can thus be considered as an ‘allosteric’ site, while the site for targeted response is the ‘active’ site of an allosteric protein. Periodic boundary condition is imposed on the side boundaries along the *x*_1_-direction.

In Fig. 2, we show the typical structure of a fit network, and the mean coordination and squared displacement maps. In the fit network, the displacements at the response site are found to be close to the expected values. The mean coordination map indicates the presence of a less coordinated region connecting the stimuli and response sites, which is surrounded by two comparatively better connected regions. The shape of this ‘floppy’ region is similar to a ‘trumpet’ with the narrow end connecting the stimuli site and the wide end connecting the response site, as observed earlier in reference [5]. This observation indicates the possible presence of allosteric chains - highly deformable or ‘liquid-like’ regions in allosteric proteins whose orientation, geometry and fluctuations are tuned to the desired functionality of the protein.

**FIG. 2.**
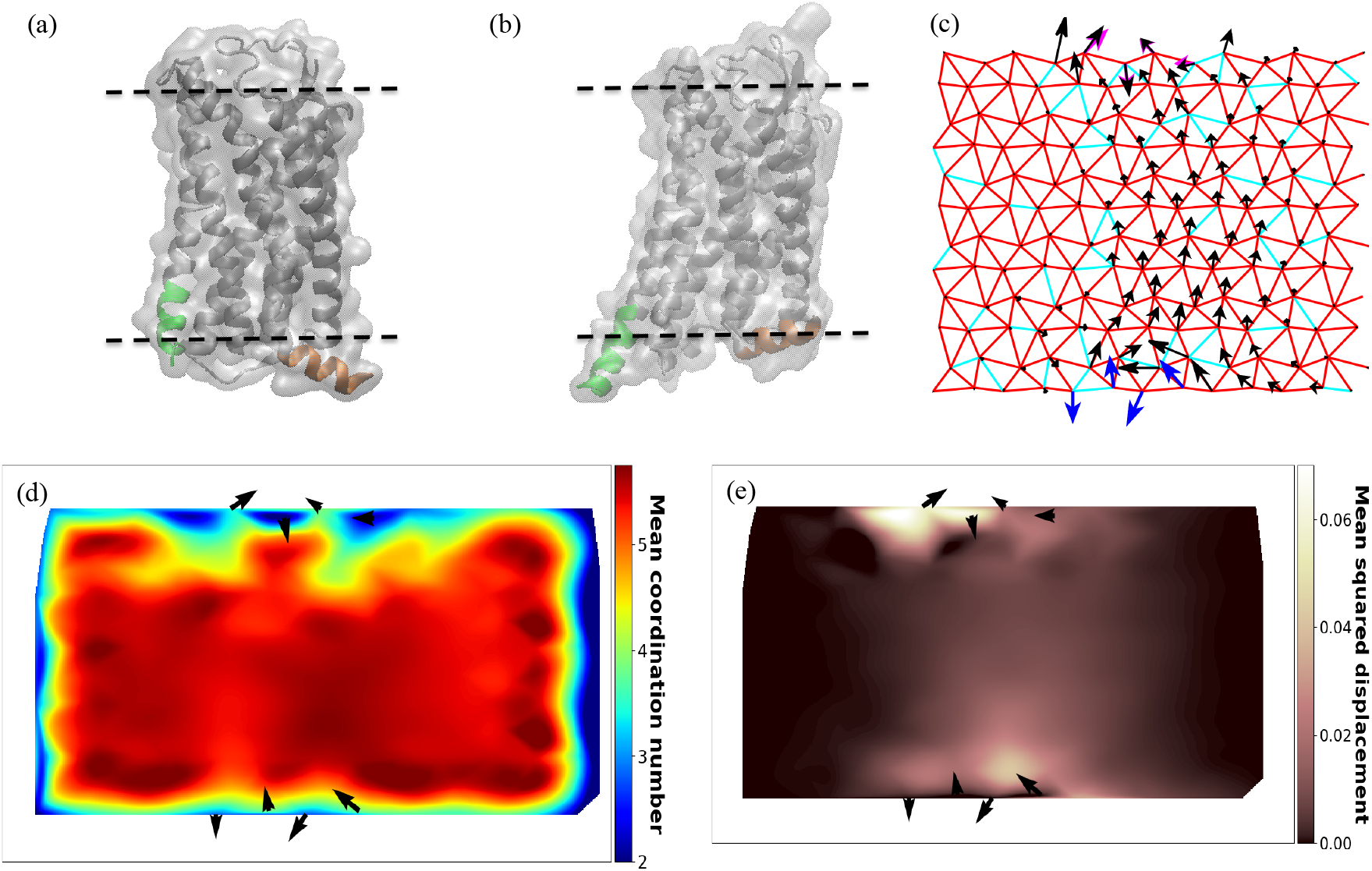
(a) The closed antagonist-bound inactive state conformation (PDB ID: 4YAY) and the (b) open agonist-bound fully active state conformation (PDB ID: 6DO1) of a GPCR protein [39] (c) An evolved optimized network. Red and cyan in the network indicate strong and zero bonds, respectively. Blue arrows indicate the imposed stimuli at the allosteric site (4 nodes at the bottom boundary), Magenta arrows on the top boundary indicate the expected response at the active sites, and black arrows are the the response field of the optimized network. (d) Average coordination number map and (e) Mean squared displacement map of the optimized network.

In a strained elastic network, the deformations die down fast away from the site of the applied strain. However, in this case the deformations, measured in terms of the mean squared displacements at all the vertices of the network, decrease far away from the stimuli sites and peak again near the response sites. This feature is also noticed for the fit abstract networks in all the other cases considered. Such an observation again indicates the presence of highly deformable regions in the protein, which can allow the strain to propagate.

Implications for structure of potentially allosteric proteins are oligomers resulting from the assembly of proteomers associated in such a way that the molecule possesses at least one axis of symmetry. The oligomeric structure creates a potentially cooperative assembly of subunits (as noted by MWC). It remains to be seen from a detailed finite size analysis whether this continuous pathway of soft interaction from I to O will be retained when we increase the size of the protein.

### B. Hinge behavior commonly found in kinases

Proteins such as Adenylate Kinase (ADK) and Guanylate Kinase undergo open-to-closed state structural transition, in order to perform their catalytic action. We model such kind of conformational change in our abstract model by fixing the response sites at the top boundary of the network, where half of the vertices have expected displacements that are rotated w.r.t. the other half. Through this, we intend to model the open-close motion of multi-domain proteins, such as ADK. The other two boundaries along the *x*_1_-direction are kept open with no periodic boundary condition.

In Fig. 3, we show the structure of a fit network and the mean coordination and squared displacement maps. The fit network is observed to be divided into two very rigid domains by a weakly connected liquid-like region, that connects the stimuli and response sites. The two rigid domains are weakly connected near the allosteric (stimuli) sites, which mimics the hinge region of the kinases around which, the rigid domains opens and closes (Fig. 3a, 3b).

**FIG. 3.**
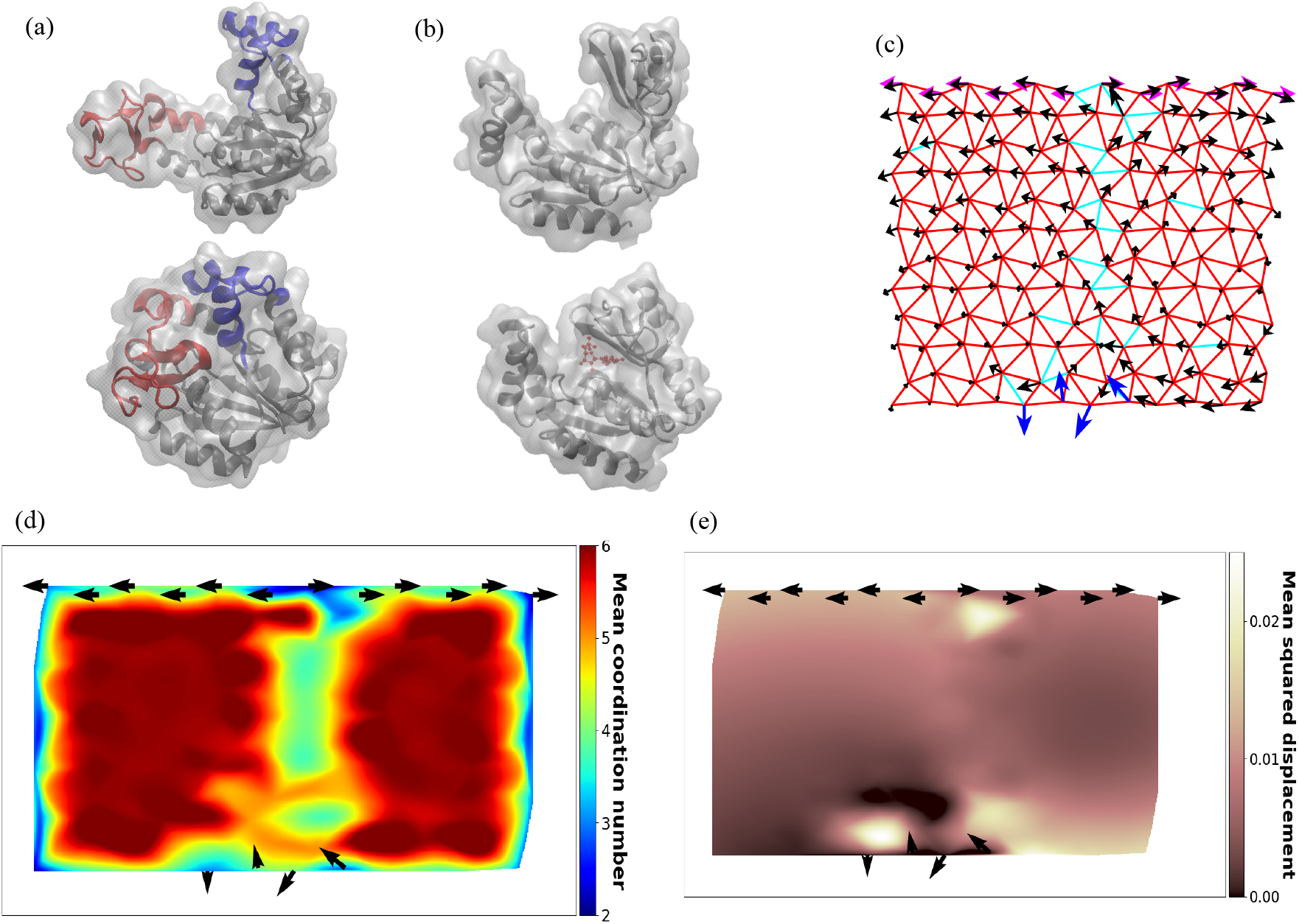
(a) The open-state conformation [PDB ID: 4AKE](top) and the closed state conformation [PDB ID: 1EX6] of Adenylate Kinase [21]. (b) The open-state conformation [PDB ID: 1EX6] (top) and the ligand-bound closed-state conformation [PDB ID: 1EX7] of Guanylate Kinase [23]. (c) An evolved optimized network with all the top boundary nodes as active sites. Red and cyan in the network indicate strong and zero bonds, respectively. Blue arrows indicate the imposed stimuli at the allosteric site (4 nodes at the bottom boundary), Magenta arrows on the top boundary indicate the expected response at the active sites, and black arrows are the response field of the optimized network. (d) Average coordination number map and (e) Mean squared displacement map of the optimized network.

### C. Conformation changes due to “buried” active sites becoming solvent exposed

In this case, we intend to model the subsequent exposure of buried residues upon ligand binding at the target sites, such as in case of GTPase, Maltose binding protein (MBP) and Calmodulin. We do this by fixing the response site at 4 consecutive vertices in the bulk of the network, with target displacements perpendicular to the bottom boundary. Periodic boundary condition is imposed along the *x*_1_-direction as in case-A, for globular allosteric proteins.

Fig. 4, shows a fit network and the mean coordination and squared displacement maps. The mean coordination map in this case is seen to be very different from the earlier two cases.

**FIG. 4.**
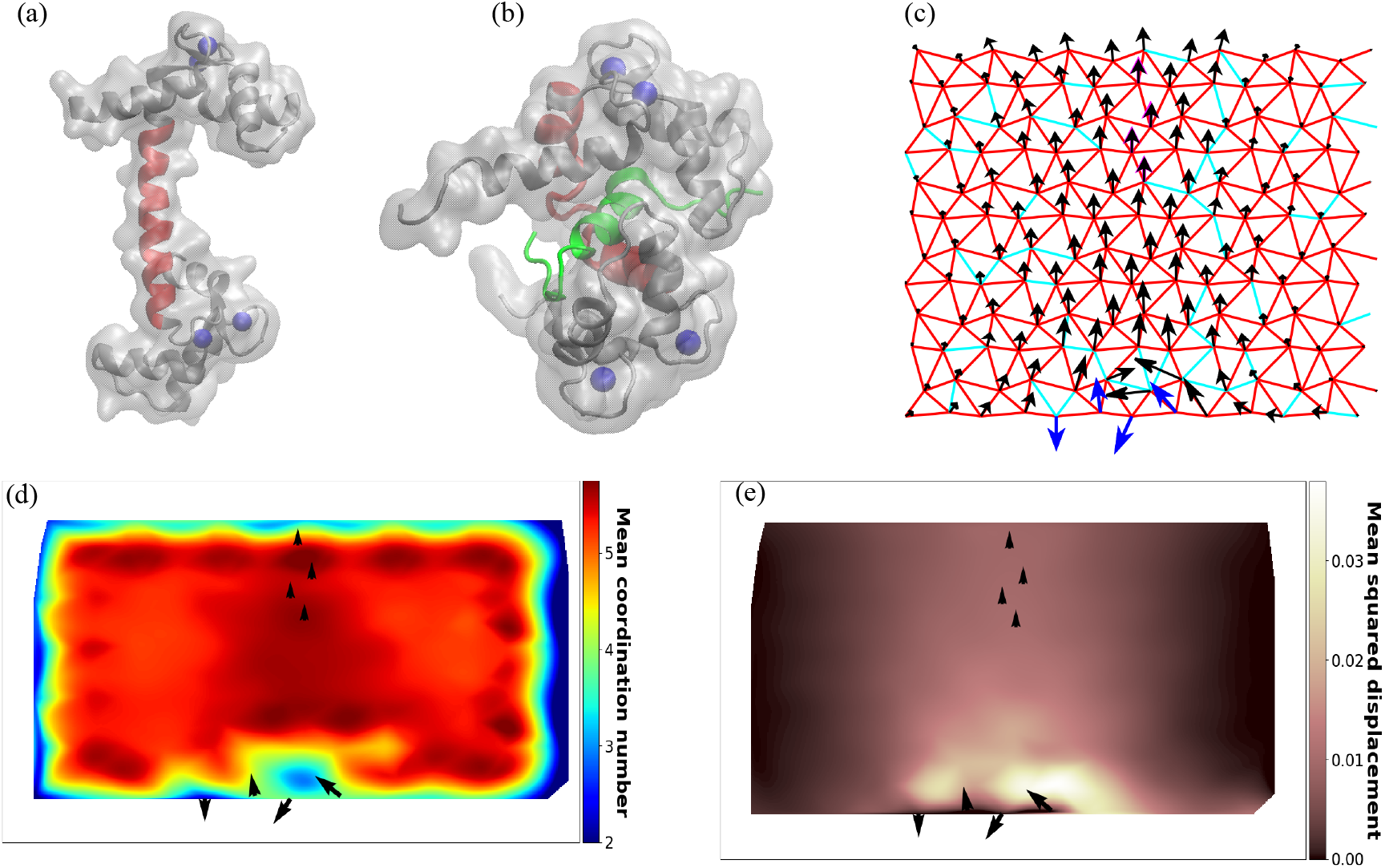
(a) The open-state conformation of Calmodulin (PDB ID: 3CLN) and (b) the peptide-bound state conformation (PDB ID: 1CKK) [27, 28]. (c) An evolved optimized network with 4 nodes in the bulk as active sites. Red and cyan in the network indicate strong and zero bonds, respectively. Blue arrows indicate the imposed stimuli at the allosteric site (4 nodes at the bottom boundary), Magenta arrows in the bulk indicate the expected response at the active sites, and black arrows are the the response field of the optimized network. (d) Average coordination number map and (e) Mean squared displacement map of the optimized network.

The response site is located within a strongly connected region, with a weakly coordinated region around it. This liquid-like region surrounds the response region on both sides and is connected at the site of stimuli. One can think of the rigid response region as the calcium binding sites of Calmodulin that stay on the rigid surface of the protein, while the low connected regions are the two target sites that open up when calcium is bound.

### D. Hinge and twist motion as in chapereone proteins

Molecular chaperone like HSP90 undergo open-to-closed state structural transition that involve large domain movements. Here we model such functional proteins in terms of the abstract network, where the response site consists of the two side boundaries with target displacement that are rotated with respect to each other. Through this representation, we try to model the hinge motions of proteins consisting of two distinct domains. As the boundaries along *x*_1_-direction serve as the response sites, no periodic boundary condition is applied in this case.

Fig. 5, shows the structure of a fit network and the mean coordination and squared displacement maps. The displacements at the two boundaries of the fit network are found to be very close to the expected response. The mean coordination map indicates a very weakly connected region in the middle of the network, similar to that observed in Case-B. However, unlike the former, the liquid-like region does not connect the stimuli and response sites. Rather, the network is divided into two very rigid domains, which move in opposite directions. As in Case-B, the liquid-like region is connected at the site of applied stimuli, which acts like the hinge region. In terms of the HSP90 example, the two rigidly connected regions can be thought of as the two flexing arms, which render the open and close form of the protein (Fig. 5a, 5b).

**FIG. 5.**
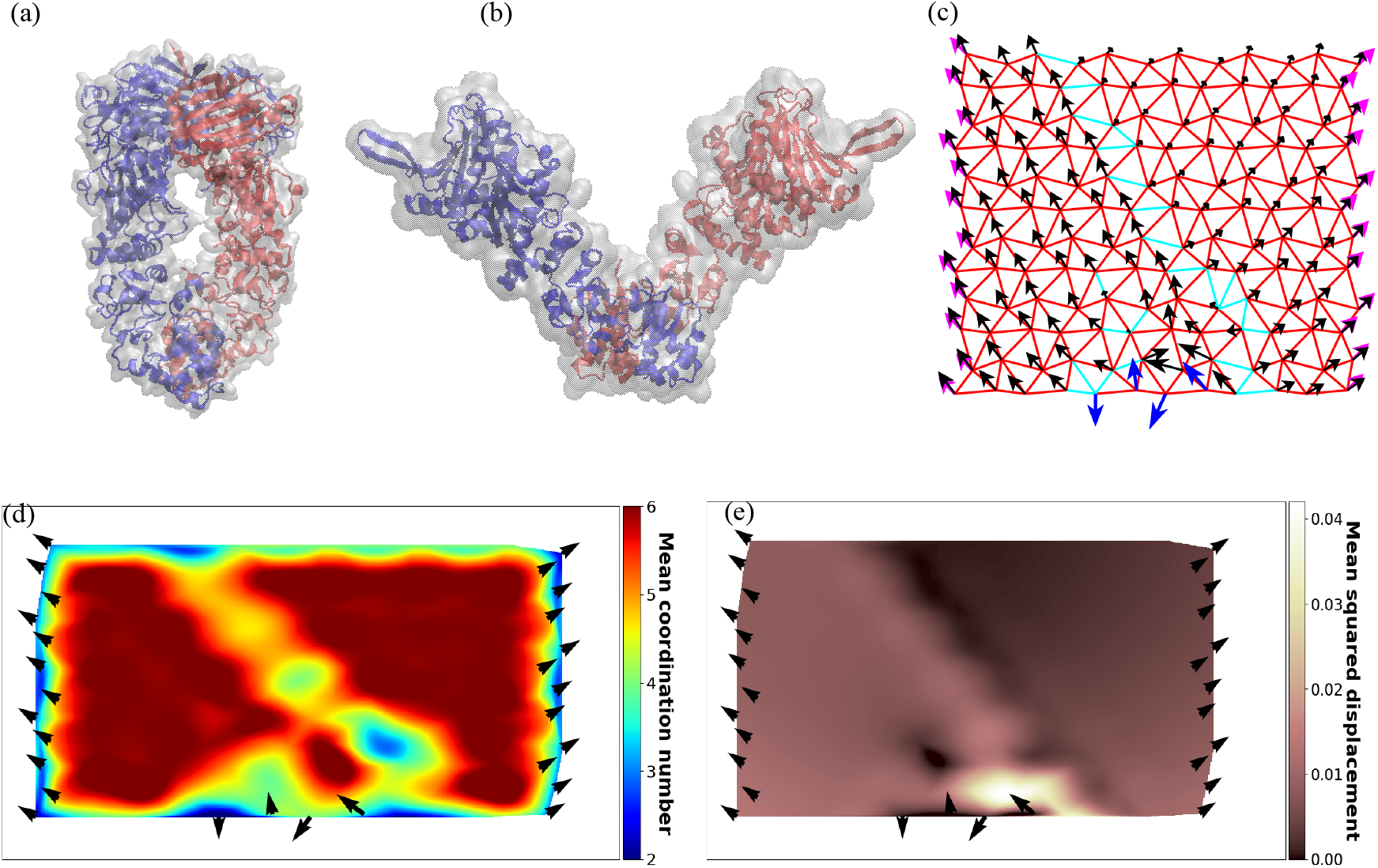
(a) The closed-state conformation of HSP70 (PDB ID: 2IOP)and (b) the open active state conformation (PDB ID: 2IOQ) [26]. (c) An evolved optimized network with all nodes on the side boundaries as the active sites. Red and cyan in the network indicate strong and zero bonds, respectively. Blue arrows indicate the imposed stimuli at the allosteric site (4 nodes at the bottom boundary), Magenta arrows on the side boundaries indicate the expected response at the active sites, and black arrows are the the response field of the optimized network. (d) Average coordination number map and (e) Mean squared displacement map of the optimized network.

## III. LOCALIZED SOFT CHANNELS AND NON-AFFINE ELASTICITY

The measured quantities evaluated on the configuration or graph that optimizes the fitness has distinct features in each of the examples studied. Each of them have a contiguous channel comprising vertices with low coordination number (relatively low constrained vertices) and large displacements, sharply separated from regions with high coordination number (highly constrained vertices) and low displacements. When embedded in a bath of temperature *T*, these low coordination number channels will be associated with large volume fluctuations, such volume fluctuations have been observed to accompany structural changes along allosteric paths [40]. The channels resemble a liquid channel embedded in an amorphous solid, and exhibit a distinct geometry and orientation.

To proceed with this intuition, we first note from Eq. 1, that the optimal configurations are minimisers of the “energy”, 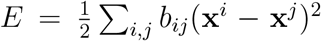, subject to constraints implied by the fixed input and desired output. These constraints can be either taken to be hard constraint, in which case these vertices are pinned, or soft constraints, represented as a term in the energy that represents the fitness function. This harmonic energy *E* can be formally represented through the spectral properties of the graph Laplacian *L* [41]. The graph Laplacian *L* acts on functions defined on the graph 𝒢. Let *u* be a real-valued function on 𝒢, i.e., *u* : 𝒱 → ℝ, with inner product

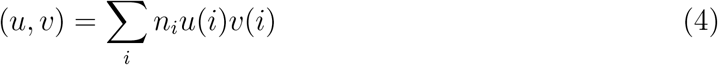

where *n*_*i*_ is the degree of *i*. Consider an operator Δ on this space of functions whose action on function *u* is

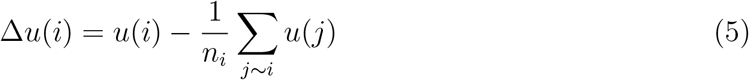

If *g* is an arbitrary function on 𝒢 (and therefore, one can view *g* as a column vector), then

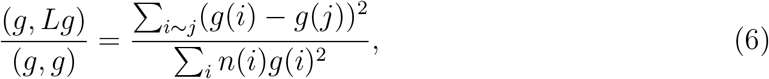

which will clearly highlight the interface of the liquid-solid regions. The spectrum of the graph laplacian describes the interface fluctuations. One can study the evolution of the eigenvalues and eigenvectors of *L* as one moves through the genotype space towards the optimal configuration.

To this graph Laplacian we add the constraints implied by the fixed input and desired output. The corresponding “Hamiltonian” graph operator that acts on functions on the graph, is described by an elliptical operator of the form *L*_*G*_ + *V*, where *L*_*G*_ is the graphLaplacian on the network *G* and *V* is the potential that imposes this constraint in 𝒫. A simple choice for *V* in Sect. II A is

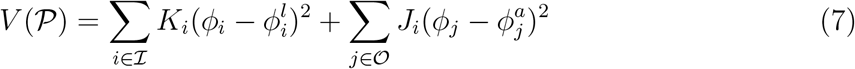

where *ϕ* is the scalar function defined on *G* (e.g., local coordination number (density) or root-square displacement) and the coefficients *K*_*i*_, *J*_*k*_ are large so as to impose the constraint strongly. This acts like a pinning potential in the target space of ℐ and 𝒪.

The Hamiltonian we have constructed bears a close resemblance to the Cahn-Hilliard theory describing the fluctuation spectrum of a pinned liquid-gas interface,

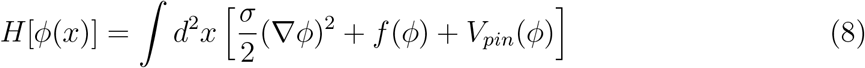

the last term is a pinning potential that breaks the Euclidean invariance of the interface [42].

The lowest eigenvalues of this model [42] includes a capillary and peristaltic mode, which resembles the liquid-like excitations of the channel shown in Fig. 2.

Another perspective is from the theory of amorphous solids. One may think of the elastic network as a realisation of an amorphous solid, and ask how one may systematically tune the properties of the amorphous solid so as to get the desired phenotype [43, 44]. The “energy”, 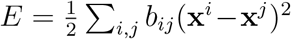, is equivalent to an elastic energy functional ∫_*x*_ ℬ (*x*)(∇*u*)^2^, where *u* is the local displacement field and ℬ are the local elastic moduli. With B taken to be randomly distributed about a mean, this is equivalent to the non-affine elastic theory of amorphous solids [45]. Now starting with a network where all the bonds are stiff, one imposes the local stress and response displacements at ℐ and 𝒪. All the bonds in the network will then undergo deformation resulting in a high elastic energy. We then make the stiffnesses of the most deformed bonds weaker ensuring that the constraints at ℐ and 𝒪 are maintained - this results in a lowering of the energy. The network obtained as a result of this “pruning” [44] will be the optimal network described above. This procedure corresponds to a random annealing of the elastic moduli to arrive at the optimal protein. The optimal solution arrived at in the example of the allosteric protein is akin to shear-banding in amorphous solids [46].

## IV. DISCUSSION

In this paper we explore ideas around a functional protein as an amorphous solid, designed to perform a specific function with high fidelity. The examples we have studied include proteins that exhibit allosteric changes such as hinge joint (e.g., Adenylate kinase and HSP90), crack formation (e.g., Calmodulin) and slide bolt (e.g., GPCR). Here we have explored the mechanical rather than the chemical facets of such a mechano-chemical machine.

This mechanical approach highlights some general points of principle. For instance, it is generally believed that in the native state, the packing density is high, making it too restricted to exhibit the variety of ways in which allostery manifests. Our analysis suggests that the native state should be allowed to be locally compressible (looser packing) thus exploring a higher dimensional low energy landscape.

Our results should remind us of the concept of *sectors* [47], envisaged as evolutionarily conserved, spatially organized molecular motifs that can enable perturbations at specific surface positions to rapidly initiate conformational control over protein function.

The optimization of fitness ℱ over the space of phenotypes is not convex, implying that there will be many solutions to the optimisation problem. In future work, we will study the geometry of the fitness landscape, the number of minima and maxima and their proximity to one another. If there are a small number of optimal solutions then one might expect these optimal features have been arrived at multiple times in the evolutionary history of proteins, thereby explaining the frequent reemergence of protein architectural motifs.

Many extensions of this work can be envisaged, such as extension to 3D, separating the backbone covalent interactions from the rest of the interactions, and including nematic correlations representing the effect of secondary structures [48]. We hope to take up these questions in the future.

## ACKNOWLEDGEMENTS

It is a pleasure to present this article as part of Prof. Somdatta Sinha’s Festschrift. At a time when Theoretical Biology was not very popular in India, it was scientists like Somdatta who bravely kept the intellectual flame burning. Somdatta continues to be a gracious mentor to the younger generation of biophysicists.

There is yet another reason for us to thank Somdatta. While the ideas presented here have been brewing with us for a very long time, we have, for a variety of reasons, not been able to present them formally. This celebration of Somdatta’s contribution to Theoretical Biology in India provides an ideal occasion to do so.

MR acknowledges support from the Department of Atomic Energy (India), under project no. RTI4006, and the Simons Foundation (Grant No. 287975). MR and SS acknowledge the award of JC Bose Fellowships, JCB/2018/000030 and JBR/2020/000015, respectively, from SERB-DST, India. A.S. thanks the Department of Science and Technology of India for the early career reward (ECR) grant. MT acknowledges the research fellowship from the Department of Biotechnology, India. This research was also supported by the Department of Biotechnology, Government of India in the form of IISc-DBT partnership programme.

## Notes

https://github.com/codesrivastavalab/allostery-theory

### Competing Interest Statement

The authors have declared no competing interest.

